# Multimodal fusion of multiple rest fMRI networks and MRI gray matter via multilink joint ICA reveals highly significant function/structure coupling in Alzheimer’s disease

**DOI:** 10.1101/2023.02.28.530458

**Authors:** K M Ibrahim Khalilullah, Oktay Agcaoglu, Jing Sui, Tülay Adali, Marlena Duda, Vince D. Calhoun

**Affiliations:** Tri-institutional Center for Translational Research in Neuroimaging and Data Science (TReNDS), Georgia State University, Georgia Institute of Technology, Emory University, 55 Park Place, NE, 18th floor, Atlanta, Georgia, USA, 30303; State Key Laboratory of Cognitive Neuroscience and Learning, Beijing Normal University, Beijing, China; Department of Electrical and Computer Engineering University of Maryland, Baltimore County, Baltimore, MD, 21250, USA

**Keywords:** Parallel Multilink jICA, Multiple fMRI networks, Multimodal Fusion, Posterior Cingulate, Parahippocampus, Thalamus, Overlapping fMRI, AD

## Abstract

In this paper we focus on estimating the joint relationship between structural MRI (sMRI) gray matter (GM) and multiple functional MRI (fMRI) intrinsic connectivity networks (ICN) using a novel approach called multi-link joint independent component analysis (ml-jICA). The proposed model offers several improvements over the existing joint independent component analysis (jICA) model. We assume a shared mixing matrix for both the sMRI and fMRI modalities, while allowing for different mixing matrices linking the sMRI data to the different ICNs. We introduce the model and then apply this approach to study the differences in resting fMRI and sMRI data from patients with Alzheimer’s disease (AD) versus controls. The results yield significant differences with large effect sizes that include regions in overlapping portions of default mode network, and also hippocampus and thalamus. Importantly, we identify two joint components with partially overlapping regions which show opposite effects for Alzheimer’s disease versus controls, but were able to be separated due to being linked to distinct functional and structural patterns. This highlights the unique strength of our approach and multimodal fusion approaches generally in revealing potentially biomarkers of brain disorders that would likely be missed by a unimodal approach. These results represent the first work linking multiple fMRI ICNs to gray matter components within a multimodal data fusion model and challenges the typical view that brain structure is more sensitive to AD than fMRI.

## Introduction

Various noninvasive neuroimaging techniques have provided remarkable new insights into human brain mapping to investigate the structural and functional changes of the human brain during transition from healthy aging to psychiatric disease. In the context of the brain imaging studies, there are different imaging techniques to acquire neuroimaging data for visualizing neural activity, improving understanding of brain mechanisms, and identifying biomarkers-especially for mental disorder. A large number of studies collect multiple modality data such as structural magnetic resonance imaging (sMRI), functional magnetic resonance imaging (fMRI), electroencephalography (EEG), diffusion tensor imaging (DTI), magnetoencephalography (MEG), and other types of data from the same individual. Among them, resting-state functional magnetic resonance imaging (rs-fMRI) has been progressively applied to functional mapping. It is increasingly evident that changing functional activity is a reliable indicator of brain disorders for early diagnosis and treatments (Whitfield-Gabrieli et al., 2009; Li, A. et al., 2020). Independent component analysis (ICA)-based mapping has been widely used, as no a prior information is required, and thus can be used to explore the entire dataset. ICA performs independent source separation and noise component removal simultaneously (Du, Y. et al., 2016a). ICA-based functional mapping is a data driven method, which is capable of identifying functional networks while preserving more single-subject variability (Yu, Q.B. et al., 2017). However, components from different subjects using ICA might not have spatial correspondence. To overcome the limitation, several methods have been proposed to obtain subject specific independent components (ICs), which preserved independence of ICs at the subject level and/or established correspondence of ICs across subjects (Du, Y. et al., 2013; Calhoun, V.D. and de Lacy, N., 2017). Lin et. al. (2010) presented a semiblind spatial ICA approach to improve ICA performance in fMRI analysis using spatial constraints. The study of brain networks for biomarker development is becoming increasingly common. Multiple brain networks are often estimated using fMRI to quantify similar functional systems (Calhoun, V.D. and Adali, T., 2012; Marusak, H.A. et al., 2017; Van Den Heuvel, M.P. and Pol, H.E.H, 2010; Yuhui Du et al., 2020). Many approaches have concentrated on a single network of the brain instead of multiple networks to identify indicator of brain disorders in neurological and psychiatric conditions such as Alzheimer’s disease and schizophrenia (Toussaint PJ et al., 2014; Du Y et al., 2016; Chaovalitwongse WA et al. 2017), which might fail to uncover more comprehensive descriptions of altered brain connectivity than multiple networks. Therefore, the goal of this study is to fuse multiple networks and multiple modality data per subject to take maximal advantage of the join-information of the existing data.

Every imaging technique provides unique, filtered and interrelated information of brain’s structural and functional organization (V. D. Calhoun and J. Sui, 2016; J. Sui et al., 2012). Combining modalities can help to provide a more complete understanding of the human brain and its disorder as compared to unimodal studies (V. D. Calhoun and J. Sui, 2016; Sendi MSE et al., 2021). Utilizing multiple modality can leverage the joint or complementary information between them. Two categories of approaches to capture joint information from multiple data sets include univariate approaches (e.g., correlation) and multivariate approaches (e.g., ICA). Multivariate approaches focus on interrelated patterns rather than unrelated points. They can estimate all the variables jointly by working with patterns instead of just paired relationships (Purcell SM et al., 2009; Le Floch E et al., 2012; Hardoon DR et al., 2009; V. D. Calhoun et al., 2009; Vounou M et al., 2010; Liu J and V. D. Calhoun, 2014; Handwerker DA et al., 2012). Multivariate approaches can also discover hidden factors (sources or features) that underlie sets of random variables, measurements, or signals. ICA, as one widely used approach, assumes that the input mixed signal is a linear mixture of the sources, which are statistically independent and (mostly) non-Gaussian. To estimate independent sources using ICA, statistical information higher than second order is needed. A variety of approaches for improving ICA algorithms have already been developed. The popular ICA algorithms that use nonlinear functions to generate higher-order statistics can be linked to maximum likelihood estimation such as Infomax (Bell AJ and Sejnowski TJ, 1995; Lee TW et al., 1999), FastICA (Hyvarinen A and Oja E, 1997). Another algorithm named joint approximate diagonalization of eigenmatrices (JADE) explicitly computes fourth order cumulants of whitened signals (Cardoso JF and Souloumiac A, 1993). Though, both FastICA and Infomax are robust, both make some assumptions about the source distributions. They work well for symmetric distributions and are less accurate for skewed or multimodal distributions and for sources, which are close to Gaussian (V. D. Calhoun et al., 2009; Koldovský Z et al., 2006; X. -L. Li and T. Adali, 2010; Boscolo RH et al., 2001; Bach F. and Jordan M., 2002; G.S. Fu et al., 2014; W. Du et al., 2016). Both algorithms require the nonlinearities to tie the form of source distribution according to a given optimality condition. There are several adaptation strategies which have been developed to relax this requirement. Koldovský Z et. al. introduced a generalized Gaussian density model (Koldovský Z et al., 2006). Other flexible adaptation methods include entropy bound minimization (EBM) (X. -L. Li and T. Adali, 2010), non-parametric ICA (Boscolo RH et al., 2001), kernel ICA (Bach F. and Jordan M., 2002) and other more recent approaches such as those based on entropy rate minimization that make use of both higher-order statistics and sample dependence (G.S. Fu et al., 2014; W. Du et al., 2016).

Badhwar et al. (2017) studied multiple regions of rs-fMRI and observed changes in functional activity with Alzheimer’s disease (AD). Zhang et al. found that the dysfunction of salience network (SAN) and default mode network (DMN) may lead to discrepancy in attention in mild cognitive impairment (MCI) and AD patients (Zhang, Y. et al., 2018). Dai et al. (2014) proposed a metric that showed abnormality regions and how they were distributed in the brain. They observed significant functional changing in DMN such as inferior parietal cortex and medial prefrontal gyrus and in some other hubs such as insula, thalamus, and supplemental motor area in patients with AD. Zheng et al. also studied rs-fMRI and showed that AD group had lower functional connectivity than healthy control (HC) especially within DMN, visual network (VN) and sensorimotor network (SMN) (Zheng, W. et al., 2017).

Alzheimer’s disease is known to impact both function and structure, but the bulk of the focus has been on structural MRI (rather than functional MRI) as this tends to show higher sensitivity and predictive accuracy. However, there has been little work focused on the identification of functional and anatomical interactions among different brain regions (Bozzali M. et al., 2011; Brier MR et al., 2014). Luo et al. (2020) showed structural-functional relationship using a dataset of over 1500 individuals, but they used a two-step process, first performing separate, rather than joint, analysis. Sendi MSE et al., 2021 studied the progression from normal brain to very mild Alzheimer disease (vmAD) using separate multimodal analysis. Jointly analysis multiple types of imaging data from same individual can be particularly useful in this regard. Qi et al. (2021) developed a three-way parallel group ICA fusion method to discriminate schizophrenia and controls. Duan et al. (2020) proposed a multimodal fusion approach called aNy-way ICA, which is capable of detecting multiple linked sources over any number of modalities without involving orthogonality constrains on sources. Joint independent component analysis (jICA) is another useful data fusion approach which allows us to take advantage of the cross-information among different modalities (Moosmann M. et al., 2008).

However, jICA has a number of limitations. First, it assumes that different modalities come from the same distribution. Secondly, typically the rich fMRI data is reduced to a single map per subject, e.g., the amplitude of low frequency fluctuation (ALFF) (Hare SM et al., 2017; Turner JA et al., 2012; Zheng R et al., 2021) or a single intrinsic network like default mode. In this study, our goal was to fuse gray matter (GM) and multiple rest fMRI networks, called intrinsic connectivity networks (ICNs). However, gray matter maps have a very different distribution from ICNs derived from ICA, which are already maximally independent (Yuhui Du et al., 2020). As we show, this can then lead to a local minima problem or poor optimization in the jICA. To address these limitations, we propose a novel approach called parallel multilink jICA (pml-jICA) to fuse GM with multiple fMRI networks. We compared the proposed parallel approach with the use of ‘regular’ jICA to link the GM maps with multiple fMRI networks, called multilink jICA (ml-jICA). As we show, pml-jICA can extract relationship between brain function and brain structure from multiple rest fMRI networks. There are multiple novel aspects of this paper: 1) we study the joint relationship between resting function and structure, 2) alternating learning of the jICA parameters between modalities (rather than concatenated learning) to ensure balance across modalities and can scale to more modalities, 3) inclusion of multiple ICA component maps (resting networks) as multiple rows for each subject each linked to the same GM map, and 4), analysis and visualization of multiple loading sets for each subject. Results show significant differences between specific linked sMRI/fMRI ICNs. The proposed approach reveals some of the expected areas including hippocampus, opposite overlap in default mode network (DMN), and some subcortical areas, which are notable network for AD progression from healthy to patients.

## Materials and Methods

### Imaging Data Acquisition

In this paper, the experimental data we used are from the longitudinal Open Access Series of Imaging Studies (OASIS-3). Data were collected from several ongoing studies in the Washington University Knight Alzheimer Disease Research Center over the course of 15 years (LaMontagne P. J. et al., 2019). The dataset is a compilation of MRI and PET imaging and related clinical data for 1098 participants. It has over 2000 MR sessions including multiple structural and functional sequences. We used sMRI and resting state BOLD sequences for our analysis. In order to evaluate neural activity at rest, participants were asked to lay quietly with their eyes open while two 6 minutes resting state BOLD sequences were collected. A total of 2165 MR sessions (236 1.5T scanner and 1929 3.0T scanner) for sMRI with T1w scan type, 1691 MR sessions (2 1.5T scanner and 1689 3.0T scanner) with BOLD-resting state scan type, and other varieties of scan types were included in OASIS-3 dataset. T1w images were segmented via statistical parameter mapping (SPM12, http://www.fil.ion.ucl.ac.uk/spm/). Table-I shows a detailed description of study demographics. For each participant, the imaging data, demographics information, and clinical dementia rating (CDR) scale were used at any stage of cognitive functionality. According the scale, all participant must have CDR≤1 at the time of the clinical core assessment. If one participant gained CDR 2, they were no longer eligible for the study. Table-II represents clinical dementia rating information for the participants. Among 1098 participants, 850 participants entered with cognitively normal adults; 605 of those remained normal while 245 converted to cognitive impairment at various stages with ages ranging from 42 to 95 years. The CDR scale of the additional 248 participants was greater than zero.

**TABLE I.**
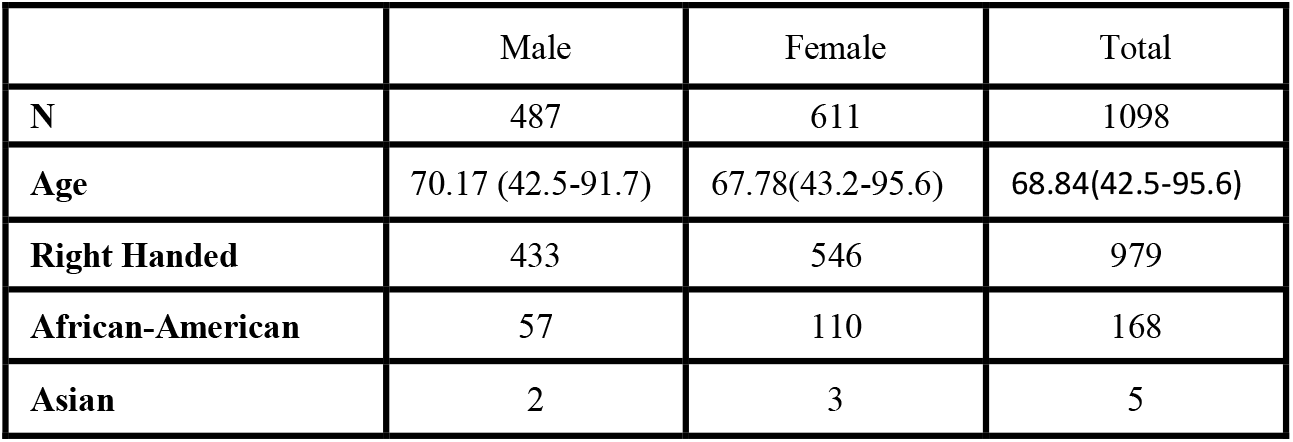
Subject Demographic Information

**TABLE II.** Clinical Dementia Ration (CDR) Distribution

|  | *Max CDR* |  |  |  |  |
| --- | --- | --- | --- | --- | --- |
| *Min CDR* | 0 | 0.5 | 1 | 2> | Total |
| **0** | 605* | 192 | 39 | 14 | 850 |
| **0.5** |  | 66 | 61 | 52 | 179 |
| **1>** |  |  | 31 | 38 | 69 |
| **Grand Total** | 605 | 258 | 131 | 100 | 1098 |

\*Healthy controls

### Quality Control and Data Preprocessing

The fMRI data were preprocessed using SPM12. Rigid body motion correction followed by slicetiming correction was performed to correct subject head motion and to account for timing difference in slice acquisition. The fMRI data were then subsequently normalized into the standards Montreal Neurological Institute (MNI) space and resliced to 3 mm x 3mm x 3 mm isotropic voxels. The resliced images were further smoothed using a Gaussian kernel with a full width at half maximum (FWHM=6mm). Next, we analyzed the data to identify 53 intrinsic connectivity networks (ICNs) using the NeuroMark pipeline, which is a fully automated spatially constrained ICA approach (Yuhui Du et al., 2020). The Neuromark_fMRI_1.0 component templates were used as spatial priors. The sMRI data were segmented and spatial normalized using the SPM unified segmentation approach, following by Gaussian smoothing with a FWHM=6mm kernel. Quality control was performed by spatially correlating each individual GM image with the group average images across all subjects and flagged scans with a correlation below 0.95. We analyzed a final set of 130 HCs and 130 AD subjects. The HCs are categorized as cognitively normal with CDR equals 0 and the ADs are categorized as AD dementia with CDR≥0.5. The age distribution of the final dataset is shown in Fig. 1. The final dataset was further preprocessed by reslicing and gaussian smoothing to match the voxel sizes (though this is not strictly a requirement of the approach). Finally, we prepared the two different types of data, GM and fMRI, for joint analysis by normalizing to unit variance across all subjects within each modality.

**Fig. 1:**
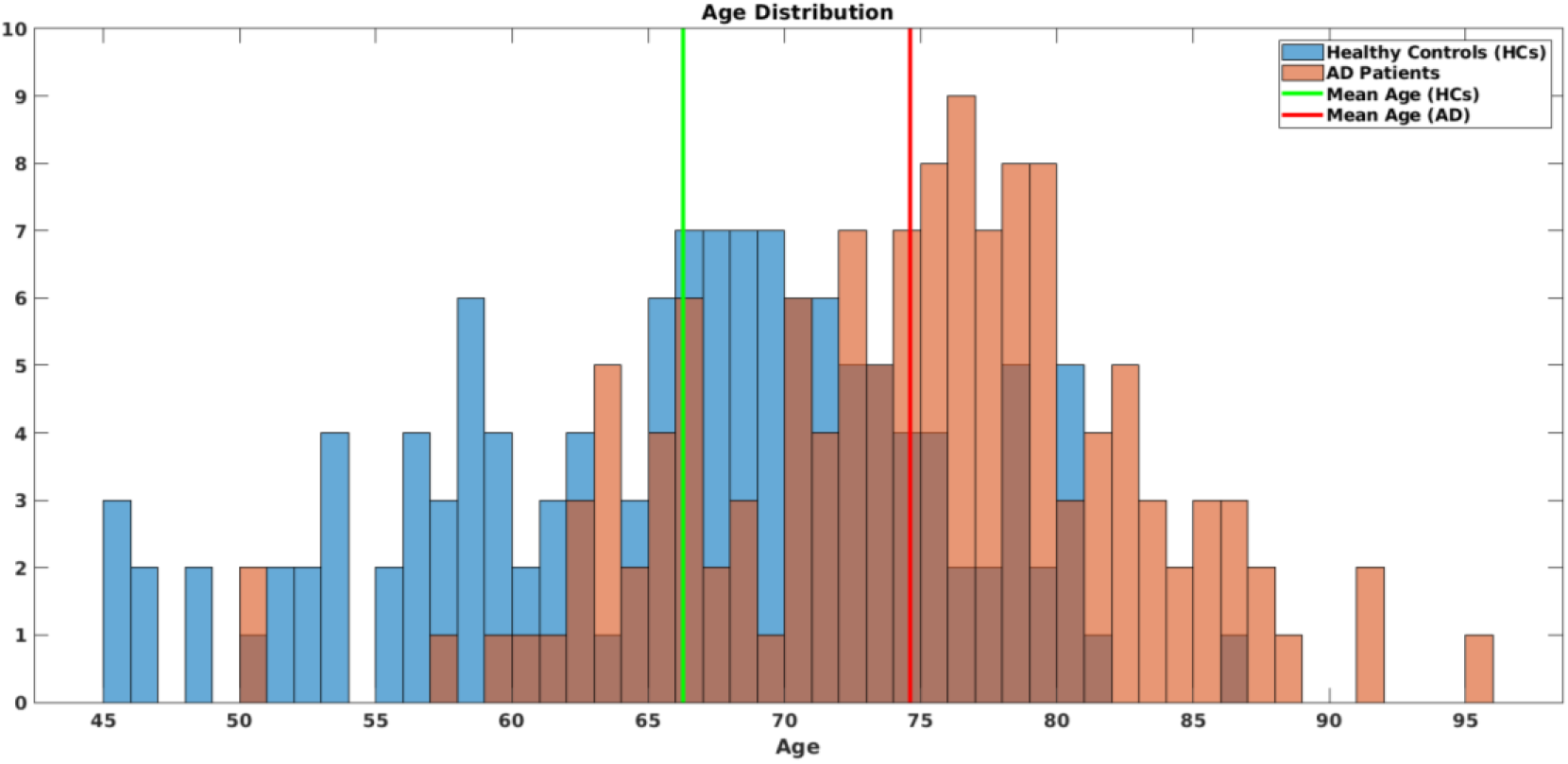
Age distribution of HCs vs ADs.

### Joint Independent Component Analysis (jICA) and multilink jICA (mljICA)

Consider a set of two equations for two sets of group data (there can be more than two, but for simplicity, we first consider two modalities):

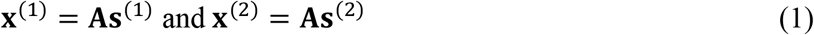

where **x**[m] and **s**[m] are the random vectors for modality m, and **A** is the mixing matrix. For a unimodal system, we can write the likelihood functions *p*(**X**^(1)^; **W**) and *p*(**X**^(2)^; **W**), considered as functions of **W**, where **W** = **A**^−**1**^, ignoring scaling and permutation ambiguities. Therefore, we maximize these two likelihood functions in a separate ICA analysis. This would give two unmixing matrices for each modality. That is, we take our estimator separately as, 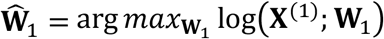 and 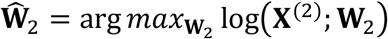. We utilize a data fusion approach to combine the two sets of optimal unmixing coefficients in the jICA.

In ml-jICA, we use the same core algorithm, but it is applied to the multiple fMRI networks through a novel approach. A conceptual model of the ml-jICA is represented in Fig. 2, which demonstrates the configuration of input feature matrix and shared mixing matrix with joint sources. Map 1 and Map 2 represent the gray matter and rest fMRI network part of the joint components, respectively. We are mainly interested in linking a targeted linear combination for multiple fMRI networks with the gray matter data. While the ICNs are maximally independent within the fMRI data, they have not been optimized for the joint fusion with GM data. We use vertical stacking of the subject specific component maps of the ICNs instead of horizontal stacking because our goal is to fuse structure and function without losing information by selecting a single ICN or reducing fMRI to a single image via ALFF or some other approaches (Hare SM et al., 2017; Turner JA et al., 2012; Zheng R et al., 2021). We do not expect the networks to covary the same way, and there are multiple ICN, so horizontal stacking was not the right solution there.

**Fig. 2:**
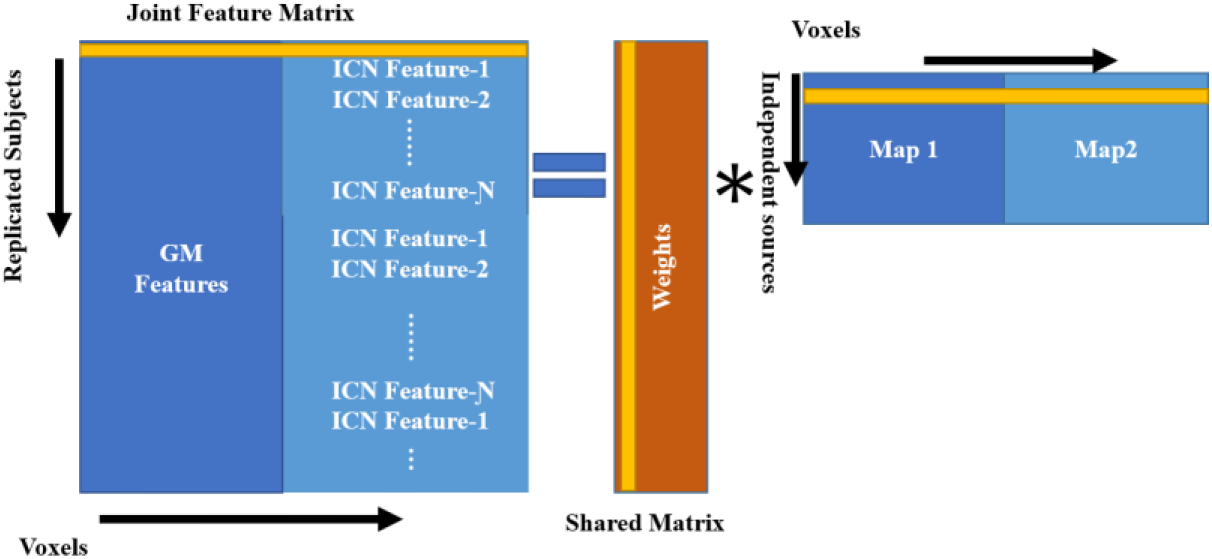
GM features and ICN features are stacked side by side to create a joint matrix. The number of ICN (Ɲ) is stacked vertically. The joint matrix is then modeled as a spatially independent joint sources images which share common mixing parameters.

In this study, to minimize memory and computation requirements, and based on prior work suggesting hippocampus and default mode as key regions of interest for AD, we investigated 8 rest fMRI networks including two ICNs which overlapped with hippocampus (48, 83) from the cognitive control domain, two ICNs including caudate nucleus (69, 99), thalamus (45), and hypothalamus (53) from the subcortical domain, and two posterior cingulate networks (71, 94) from the default mode network domain (Yuhui Du et al., 2020). These are shown in Fig. 3. The linear combination of the ICNs for a subject can be defined by the following equation,

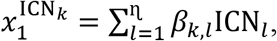

**Fig. 3:**
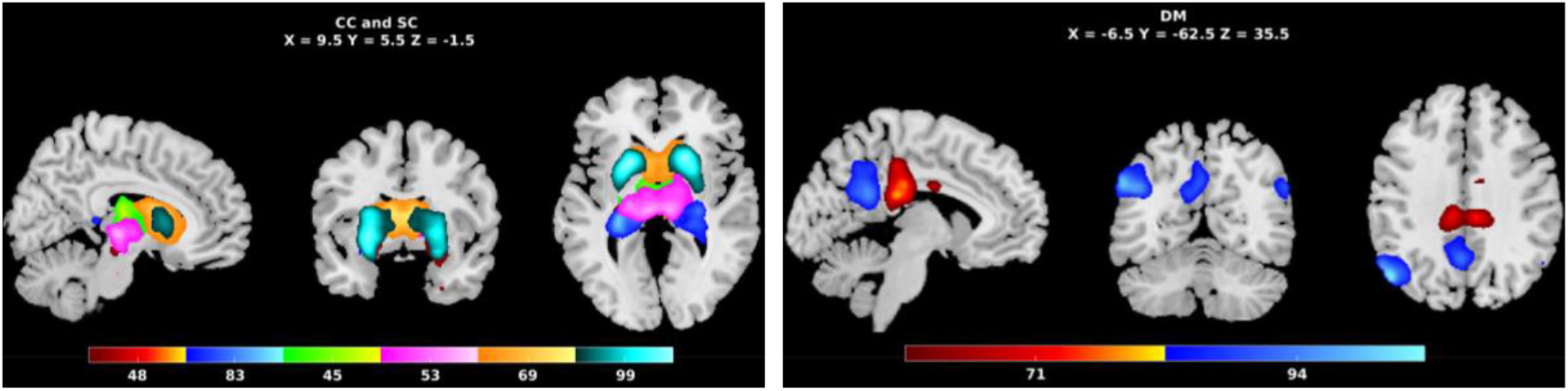
8-rest fMRI networks; CC: Cognitive Control Domain (Hippocampus (48, 83)); SC: Sub-cortical Domain (Thalamus (45), Hypothalamus (53), Caudate (69, 99)); DM: Default-mode Domain (Posterior Cingulate (71, 94)).

where 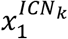 is a linear combination of the ICN for subject-1, which is replicated *k* times (*k* = 1,2, …, 8); *β*_*k,l*_ represents scaled uniformly distributed random numbers, where *l* = 1,2, . . . ɳ(ɳ = 8) for each ICN. This is applied at the data preprocessing stage before feeding into the ml-jICA and pml-jICA algorithms. Principal component analysis (PCA) is used to reduce the dimensionality of the joint multimodal input data, GM and linearly combined ICNs. Every subject specific component map of fMRI network is stacked side by side with GM. In this representation, the GM feature of each subject is replicated for the multiple rest fMRI networks.

A generative model of the given datasets, **X**^G^ and **X**^ICN^, can be written as, **x**^G^ = **As**^G^ and **x**^ICN^ = **As**^ICN^, where, for the case of two modalities and two subjects, 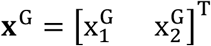 is the mixed data for the gray matter modality, and 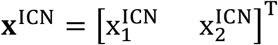 is the mixed data for the ICN (rest-fMRI modality); 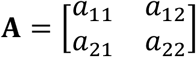 is the shared mixing coefficient matrix, and **s**^G^ and **s**^ICN^ are the gray matter and ICN sources, respectively. We can write observation vector for each subject as a single expression, 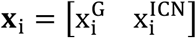, similarly for the source vector, 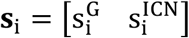; The resulting joint equation for subject *i* is then **x**_i_ = **As**_i_. The resulting joint equation for all of the subjects can be written in this matrix form,

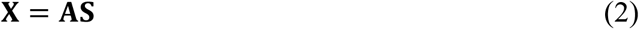

After concatenating the two datasets to form **X**^J^, the likelihood is written as,

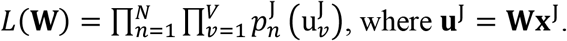

Each entry in the vectors **u**^J^ and **x**^J^ are replaced by the observation for each sample *n* = 1, …, *K*, …, *N* as rows of matrices **U**^J^ and **X**^J^. If the two sets of group data have dimensionality *N* × *V*_1_ and *N* × *V*_2_, then the joint likelihood can be written as,

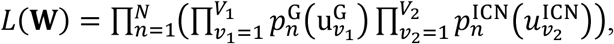

for the maximum likelihood (ML) estimator based jICA.

In this study, we utilized Infomax algorithm (Bell AJ and Sejnowski TJ et al., 1995) to estimate the common unmixing matrix for the joint analysis. The application of Infomax to source separation consists in maximizing an output entropy. It is implemented by maximizing the following ML score function with a fixed (sigmoid/tanh) nonlinear function:

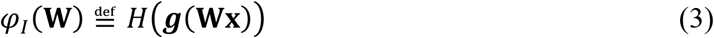

Infomax is directly linked to maximum likelihood and the nonlinearity used is a good match for super-Gaussian sources, and hence for fMRI/MRI analysis as well (T. Adali et al., 2014). For the data fusion analysis, joint data matrix is created as (we consider two modalities for simplicity), **X** = [**X**^(1)^, **X**^(2)^], placing one beside the other (see Fig. 2), similarly, the source matrix is created as **S** = [**S**^(1)^, **S**^(2)^], in which each of the rows represents the number of voxels in two images, components of the two modalities for each subject. Then, the equation (3) can be modified as,

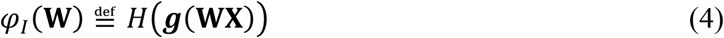

The model could jointly identify paired components by linking with multiple rest fMRI networks and optimize the decomposition for a variable of interest.

### Parallel multilink jICA (pml-jICA)

The original jICA formulation assumes that the joint sources with the two data types (GM and fMRI) modulate the same way across *N* subjects, that is, the sources have a common modulation profile across subjects as well as statistical independence among the joint sources. Assuming inter-subject variations of multimodal sources to be the same for different modalities is a strong constraint. While statistical independence may be a feasible assumption, the jICA algorithms may not be able to accomplish both of the constraints at the same time. It may end up achieving a tradeoff solution between the two constraints (Correa NM et al., 2008). In addition, the jICA model requires contributions from both modalities to be similar, so that one modality does not dominate the other in the estimation of the maximally independent components. If the two datasets have different ranges, the datasets need to be normalized in the preprocessing stage.

In this experiment, we used GM and subject specific spatial maps of the ICNs, where ICNs are already maximally independent, that is, output of ICA, which have very different distributions. Therefore, the jICA and ml-jICA models are in conflict with the i.i.d. assumptions. To address this, and generalize jICA, we proposed an adaptation approach for fusing GM and multiple rest fMRI networks by parallel optimization.

In the parallel optimization approach, the weights of each modality are optimized separately with different number of iterations for each pass. The weights are initialized for each pass by taking average after the number of iterations. Finally, the mixing loading matrix is calculated by projecting the final average weights on the joint feature matrix, which is calculated from the joint data matrix. The overall optimization approach is illustrated in Fig. 4. **X**^**1**^ and **X**^**2**^ are datamatrix for GM and fMRI with size *N* × *V*_1_ and *N* × *V*_2_, respectively, where *N* is the number of subjects and *V*_1_ and *V*_2_ are the number of voxels, and m represents the number of modalities. Each datamatrix was normalized by mean across subjects and then scaled within each modality. Before weight optimization, principal component analysis (PCA) is used to reduce dimensionality of the data, and decorrelate the data using second-order statistics, where whitening matrices, denoted as **R**_1_ and **R**_**2**_ (Fig. 4), are same. After sphering the data, the initial weights are optimized iteratively by maximizing an output entropy until ***max***_***step*** or satisfy the terminating conditions. The iteration number is controlled by ***step***_***size***. After every pass, the iterative optimized weights are denoted by **W**^G^ and **W**^ICN^ for GM and fMRI, respectively.

**Fig. 4:**
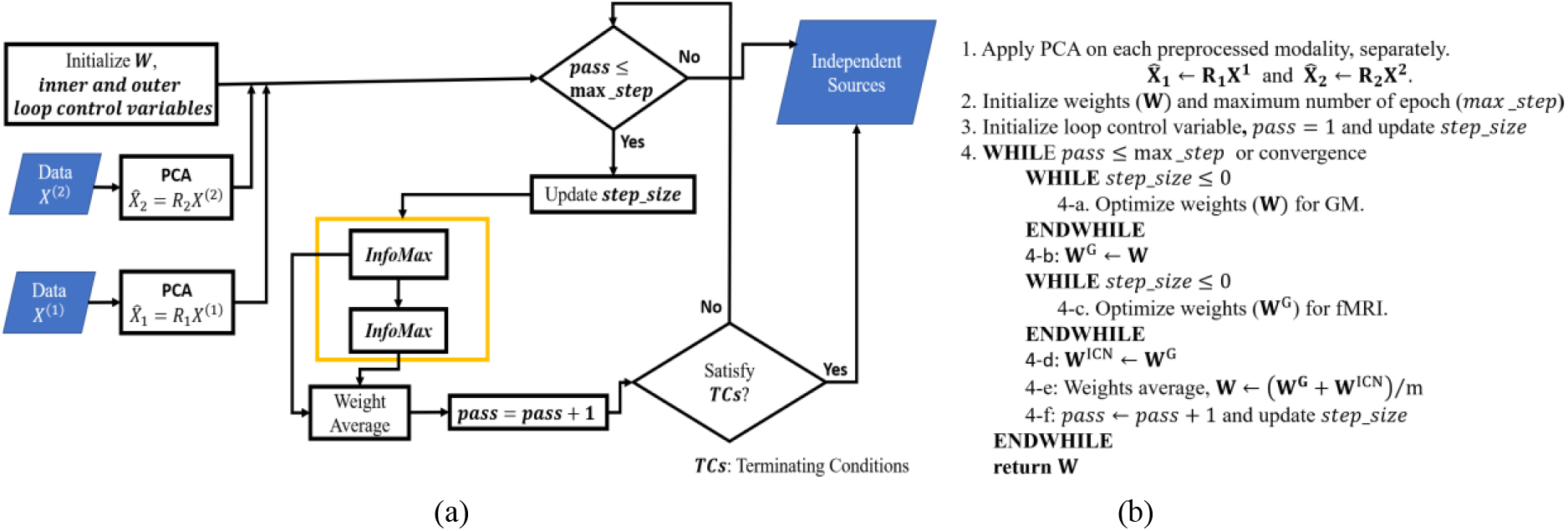
(a) Diagrammatic representation of the proposed parallel multilink jICA (pml-jICA) approach; (b) pseudo-code for the optimization approach.

## Results

In this section, we present results and performance analysis for ml-jICA and pml-jICA. Our example shows result from GM and 8 rest fMRI networks from each subject where 130 healthy controls and 130 ADs subjects are investigated in this experiment. We optimized the data matrix differently for ml-jICA and pml-jICA. The data matrix is created separately for the rest fMRI networks and GM, respectively, prior to either concatenation or joint optimization. Each subject has multiple rows corresponding to each of the 8 rest fMRI networks and the sMRI data matrix is replicated to provide corresponding columns. The number of rows for both fMRI and GM is 2080 (8 times the number of subjects) for 260 subjects. For the pml-jICA, the fMRI and GM data matrices are analyzed in a slow optimized fashion. First, the weights of the fMRI and GM are optimized with one iteration in the first pass, then optimized with four iterations from second to twentieth pass, then optimized with eight iterations from twenty-first to fortieth pass; increasing in this way the weights are optimized slowly to reach convergence. For every pass, weights are averaged before the next pass.

We estimated 15 joint components. The number of components is a free parameter, which can be either determined from empirical experience or estimated using information theoretic approaches (Beckmann CF et al., 2004; Li YO et al., 2007). The chosen component number is approximately twice of the number of networks since we were interested in a more fine-grained fusion. Based on empirical results, this number provided a good option to separate signals and noise into different independent sources. Results from the estimated components showed several interesting patterns, and also surprisingly large group differences between AD and HC for some components. The loading parameters separated by group are shown in Fig. 5(a) for the pml-jICA. The red circles and blue asterisks represent loading parameters for controls and ADs, respectively. We found several components that demonstrated highly significantly different loading parameters in patients and controls. The black line segment is the mean value of the control and the cyan line segment is the mean value of the patient. The mean difference (*min*_*diff*) between two groups show us how much they differ in patients and controls. The standard deviation of the loading parameters tells us how the weights vary across subjects.

**Fig. 5:**
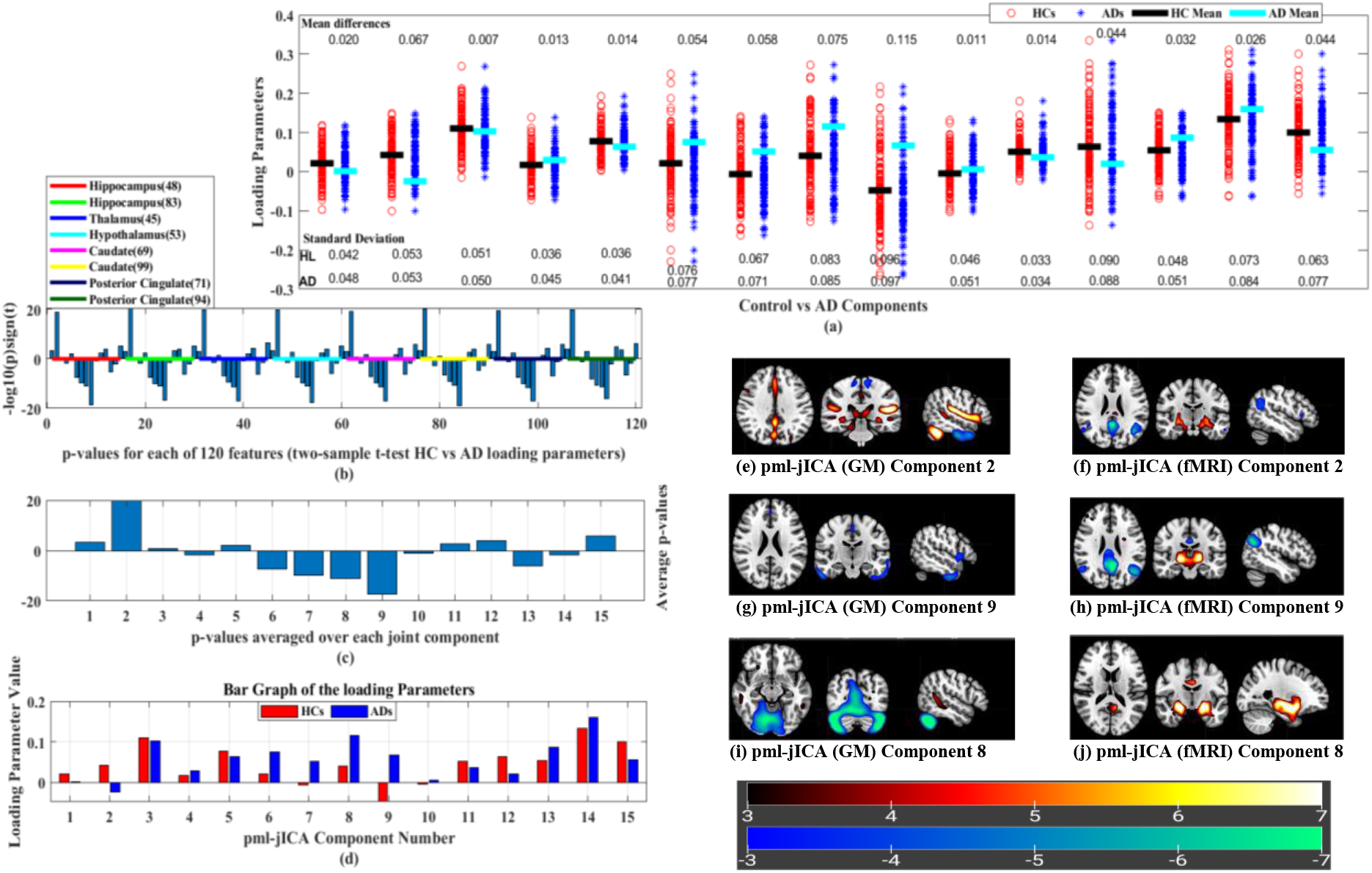
(a) Demonstrating loading parameters between Control vs *AD* components; (b) t-test between *CNs* vs *ADs*; (c) Average p-values of the rest fMRI networks; (d) Bar graph of the loading parameters between *CNs* vs *ADs*. (e)-(j) represent joint source map (pml-jICA) for components 2, 9 and 8, respectively; Splitting the overlapping fMRI DMN into two components ((f) and (h)); (e) and (f) show gray matter changes in posterior cingulate regions near the fMRI default mode posterior cingulate peak. (e)-(h) represents coupling between multiple networks including precuneus, posterior cingulate, hippocampus, and some subcortical regions. (i) and (j) represent joint relationship between cerebellar area of GM, and parahippocampal gyrus (PHG) and some regions of subcortical domain (SC) in the fMRI

To quantify correlations among the fMRI networks at rest and to find most significant components, we perform two sample t-tests between controls versus AD for each rest fMRI network with significance level 0.001. From 8 fMRI networks and 15 joint components, we estimated 120 p-values. The estimated p-values (-log10(p)*sign(T)) are shown in Fig. 5(b). The colored lines separate the p-values for each fMRI ICNs. We observe a generally similar patterns among the different networks/ICNs, though as we show later, there is also some interesting variation. After taking component-wise average of all networks, we found several significant components based on the threshold for statistical significance, which was set at corrected *P* < 0.05. The p-values for the significantly differing components 2, 6, 7, 8, 9, 13, and 15 are shown in Fig. 5(c). The average loading parameter values for the controls and patients are shown in Fig. 5(d). These show us both the degree to which the component is expressed in each group, and also whether it is positively or negatively expressed. Component 2 and 9 represent the most significant difference between controls and ADs.

The 7 joint source maps that showed significant group differences were thresholded separately for GM and fMRI. The resulting voxels shown the ones that contribute strongly to these components either positively or negatively. Multiple networks were identified for GM and rest fMRI by the proposed approach. Five of these components (6, 7, 8, 9, and 13) contained areas where joint source values were higher in ADs than controls; two components (2 and 15) had active areas where joint source values were greater in controls than patients (Fig. 5(c)). We summarized important activated brain regions of the 7 spatial patterns having volumes greater than 0.4 *cm*^3^. The details are in the supplementary materials. We described here the most interesting patterns from the selected joint independent components, especially in component 2, 8, and 9. A full summary of results is presented in the supplemental materials. The top two largest differences between diagnostic groups were found in component 2 and 9. The component 2 (*min*_*diff* = 0.067, *p* − *value* = 3.9*e* − 20) and component 9 (*min*_*diff* = 0.115, *p* − *value* = 1.82*e* − 17) formed a consistent spatial grouping, where component 2 indicates greater volume in controls than ADs and component 9 indicates lower volume in controls than ADs (Fig. 5(c)). If we notice the two components, we see that they have slightly different GM maps, and both subtractive in AD; but splitting the fMRI DMN into two different components, one is subtractive and another one is additive in AD. The most interesting finding here is that our joint analysis (pml-jICA) separates overlapping regions of the fMRI DMN into two different ICs from the input data, which is illustrated in Fig. 5(e)-(h). Another interesting finding is in the component 8, which represents joint relationship between cerebellar area of GM and parahippocampal gyrus (PHG) and subcortical domain (SC) of the brain in fMRI. Fig. 5(i)-(j) depicts spatial pattern and their activated brain regions for Component 8. Details on additional significant components (6, 7, 13, and 15) are presented in the supplemental materials.

### Model performance and comparison

Results from our comparison of ml-jICA and pml-jICA in terms of weight distribution are demonstrated in Fig. 6. This figure illustrates the weight distribution between modalities. Our expectation was that ml-jICA would be more likely to fall into a ‘modality specific’ mode due to the different distributions of GM and ICNs. We calculated the ratio of the number of strongly contributing voxels (activated voxels after thresholding) for each component between fMRI and GM, shown in Fig. 6(a)-(b). One of our observations of ml-jICA was that the learning tended to converge to either fMRI only sources, or joint sources (see Figure 6(c)-(f) for examples of this). This is likely an effect of pooling together maximally independent courses and more Gaussian-like features (in the GM maps) for the optimization, and mixing them together in a single optimization. This behavior can also be quantified by evaluating the standard deviation of the ratio, mean absolute value (MAV), and the image (Fig. 6(b)). Our pml-jICA approach was developed to learn the weights from either GM or fMRI, followed by initialization and optimization of the other modality and alternating until convergence. Visual results (Fig. (a)-(b)) showed a more balanced result where all the components are contributed to by both modalities. Results from this plot shows more extreme values for the ml-jICA, and for pml-jICA indicates that weights are more evenly contributed to by fMRI and GM than ml-jICA. Our proposed algorithm provides a solution which balances the contributions of two modalities and allows for joint estimation from multimodal data with vastly different distributions. The source maps of the two most roughly distributed components (component 7 and 13) are shown here as an illustration (Fig. 6(c)-(f)).

**Fig. 6:**
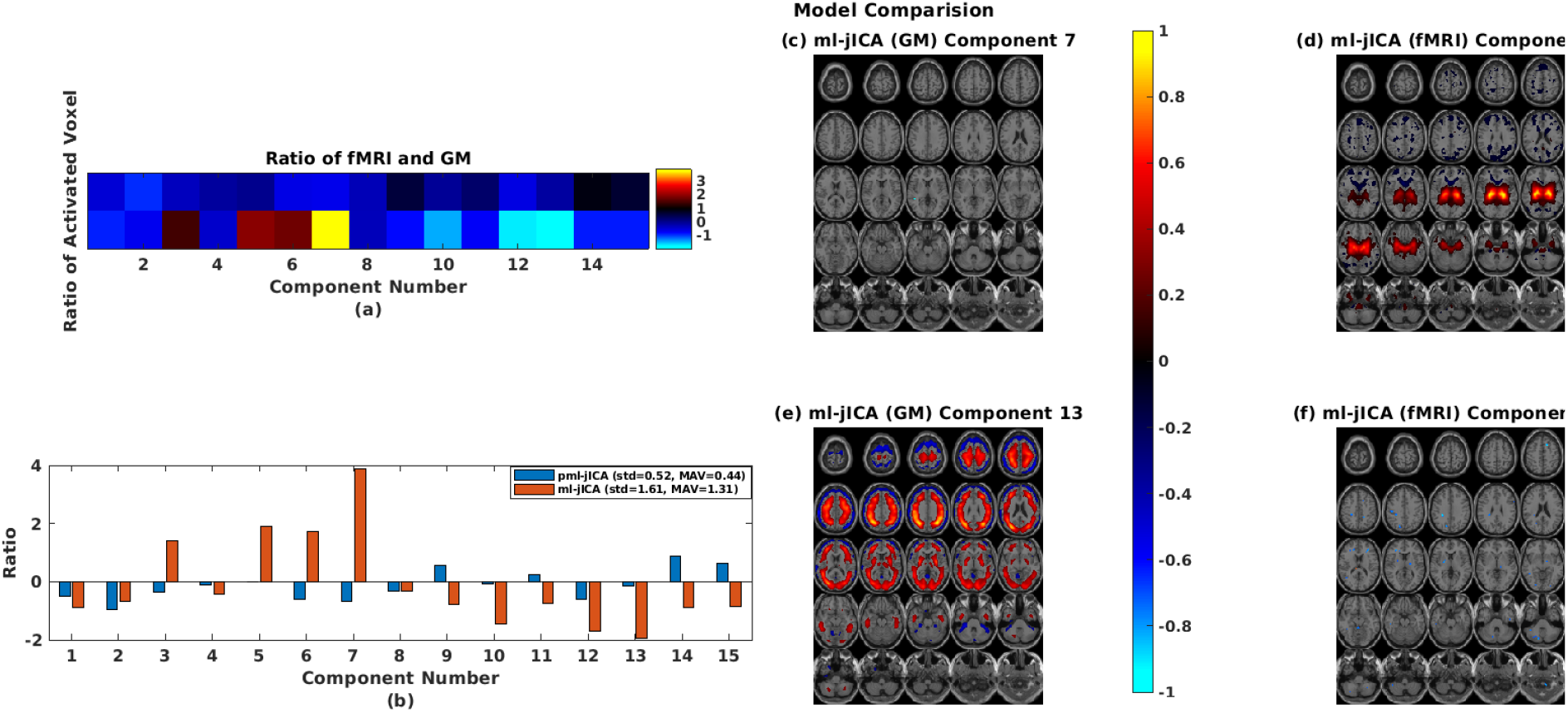
Weight distribution among modalities for model comparison. The color bar indicates the color mapping for the source maps.

### Multilink Network Contribution

While the overall patterns of multiple networks within subjects were similar, there were also some important differences. We can see this in Fig. 7, Fig. 7(a) shows loading contribution relative to mean for each subject across 8-rest fMRI networks. This highlights the variability that is present. To further unpack this, we can see in Fig. 7(b)-(g) the relative contribution for each fMRI network for *CNs* and *ADs* separately and also the difference between *CNs* and *AD*s. The right image of component 2 (Fig. 7(c)) indicates that the posterior cingulate of default mode network has greater representation in *CNs* than in ADs. Fig. 7(c) and (g) tell us that the posterior cingulate component has the strongest contribution among all of the 8-rest fMRI networks. The posterior cingulate cortex is a highly connected and metabolically active brain network. Our result highlight abnormal functional connectivity in the posterior cingulate network, which is consistent with, but significant expands, prior work (Zhang J et al., 2017; Mutlu J et al., 2016; Lee P-L et al., 2020; Yu E et al., 2017; Leech R et al., 2012).

**Fig. 7:**
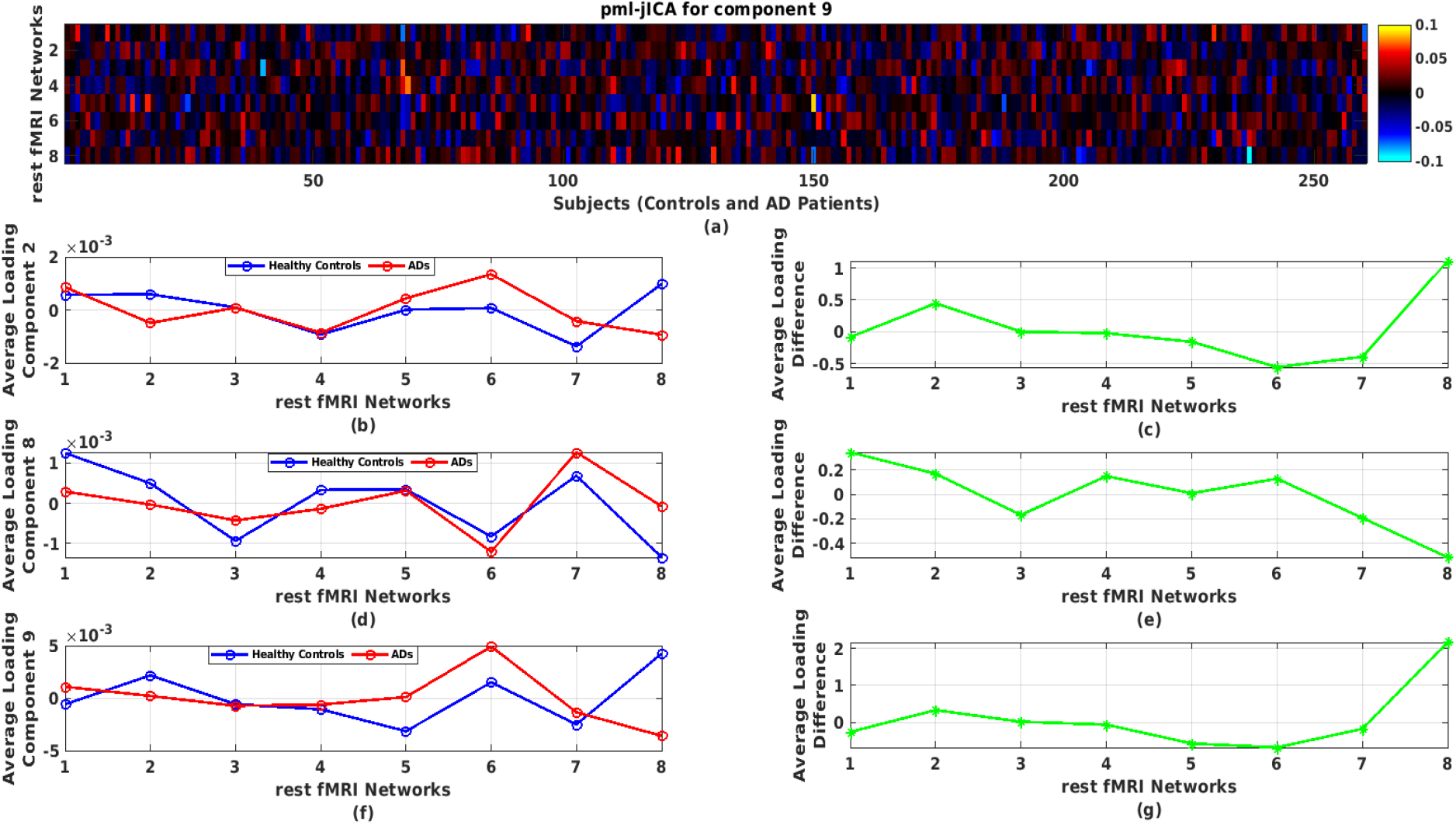
Multilink network contribution in the 2, 8, and 9 independent components; (a) representing variability in linking between different networks across subjects in the component 9; (b)-(g) demonstrating network contribution in the component 2, 8, and 9 as a representative of the 15 estimated components.

## Discussion

Our study examines the structural and functional changes between *HCs* and *AD* patients. We found interesting structural and functional pattern especially in precuneus, posterior cingulate, hippocampus, insula, superior frontal gyrus, medial and middle frontal gyrus, supplementary motor area, thalamus, putamen of the default mode, cognitive control, sub-cortical regions, respectively, as well as cerebellar and sensorimotor regions of the brain; which are known to be significant affected areas for Alzheimer’s disease (Zhang J et al., 2017; Mutlu J et al., 2016; Lee P-L et al., 2020; Yu E et al., 2017; Leech R et al., 2012; Zhong Y et al., 2014; Miao X et al., 2011; Rao YL et al., 2022; de Long LW et al., 2008; Mavroudis I., 2019; Giacomo Koch et al., 2022; W. J. Henneman et al., 2009). The DMN plays an important role in functional brain structure. The PCC has been shown to have a significant causal relation with other nodes (Miao X et al., 2011). Neuropathological atrophies are mostly distributed in posterior cortical regions, e.g. precuneus, posterior cingulate, of the brain in the early stage of AD (Mutlu J et al., 2016; Giacomo Koch et al., 2022). In addition to the DMN, the hippocampus, the area in the cognitive control (CC) of the brain that is important for memory, may be potential biomarkers to predict the progression from MCI to AD (Rao YL et al., 2022; de Long LW et al., 2008; W. J. Henneman et al., 2009). In addition, de jong LW et al. (2008) found that the volumes of putamen and thalamus were significantly changed in patients diagnosed with probable Alzheimer’s disease.

Results for the highly significant components are shown in Fig. 5(e)-(f). In the fMRI part of the joint component, angular gyrus and some parts of the DMN show negative activation whereas hippocampus shows positive activation. In the GM part of the joint component, some parts of the DMN, thalamus, parahippocampus, and cerebellum show positive activation. We also see some portions of the posterior cingulate show both fMRI and GM changes. Besides, the superior temporal gyrus is additive and inferior temporal gyrus is subtractive. Some parts of the sensorimotor (precentral, paracentral lobule) are also subtractive. The loading directionality of this component is *CNs* > *ADs* (Fig. 5(c)). So, the hippocampus represents functionally lower connectivity in AD group as well as the thalamus, parahippocampus, and cerebellar are structurally reduced in GM which consistent the previous studies (Mutlu J et al., 2016; ; Zhong Y et al., 2014; Miao X et al., 2011; de Long LW et al., 2008). In Fig. 5(g)-(h), angular gyrus and some parts of DMN (especially posterior cingulate area) have negative activation whereas thalamus has positive activation in fMRI. From GM, we see that inferior temporal gyrus of the auditory area and supplementary motor area reduce volume in *ADs* than *CNs*.

If we compare component 2 and 9 of pml-jICA, we see that our proposed approach separates overlapping regions of the DMN into the two components, which is highlights an important benefit of the pml-jICA. We observed that some portion of the DMN (especially precuneus) changes into another portion of the DMN (especially posterior cingulate) from component 2 to component 9, where one is additive and another one is subtractive, which implies that AD group has lower functional connectivity in the posterior cingulate area than healthy group. This inference is also consistent with previous studies (Mutlu J et al., 2016; Zhong Y et al., 2014; Miao X et al., 2011).

In component 8, precuneus (left), hippocampus, and putamen networks of the DMN, CC, and SC, respectively, are additive in the fMRI, whereas cerebellar network is subtractive in the GM (Fig. 5(i)-(j)). This component shows coupling between cerebellar structurally and hippocampus and some subcortical areas of the brain functionally. The hippocampus and activated subcortical areas show higher connectivity in AD group whereas cerebellar reduces in AD group.

From component 2, 8, and 9, we may infer that posterior cingulate, hippocampus, and some subcortical areas are the most important network for AD disease progression from healthy to AD patients. Our multilink network contribution analysis also tells us the same implication as the joint source analysis.

## Conclusions

We presented two multimodal fusion approaches, called ml-jICA and pml-jICA, to examine the relationship between healthy and Alzheimer’s individuals. The main purpose of this study is to discover unique components from rest fMRI networks and gray matter maps, which share similar correspondence between subjects. This then enables to identify which discovered components best differentiate between patient and control groups. There has been little work combining ICNs with GM data, and our results suggest this may ignore some information that can inform us about the impact of AD on the brain. In this initial work, we fused 8 rest fMRI networks with gray matter. The main advantage of ml-jICA approach is that it maximizes the joint likelihood function of multiple ICN or brain networks with GM providing more reasonable solution from one that does not utilize joint statistics. Also, the vertical stacking preserves more structural and functional information than horizontal stacking via ALFF or other approach. Our results represented advantages of the pml-jICA over ml-jICA in the applications of examining AD impairment. The results also suggest that posterior cingulate network is the most contributing network among the eight rest fMRI networks. In future, we will further optimize pml-jICA to incorporate more ICNs in order to provide a more complete structure/function mapping across the brain.

## ACKNOWLEDGEMENTS

This study was supported by NSF 2112455 and NIH RF1AG063153.

