## supplementary materials for "Multimodal fusion of multiple rest fMRI networks and MRI gray matter via multilink joint ICA reveals highly significant function/structure coupling in Alzheimer’s disease"

The joint source maps of the parallel multilink joint ICA (jICA) are shown in the following figure.

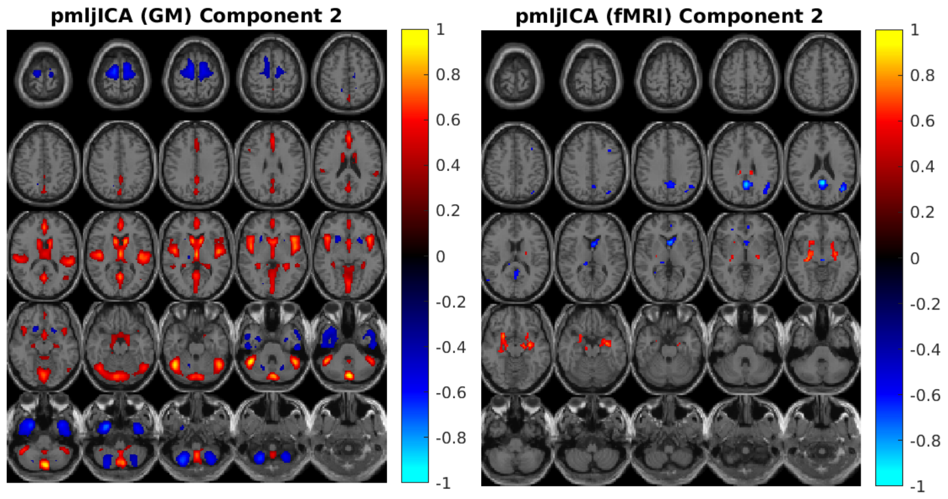


(a)

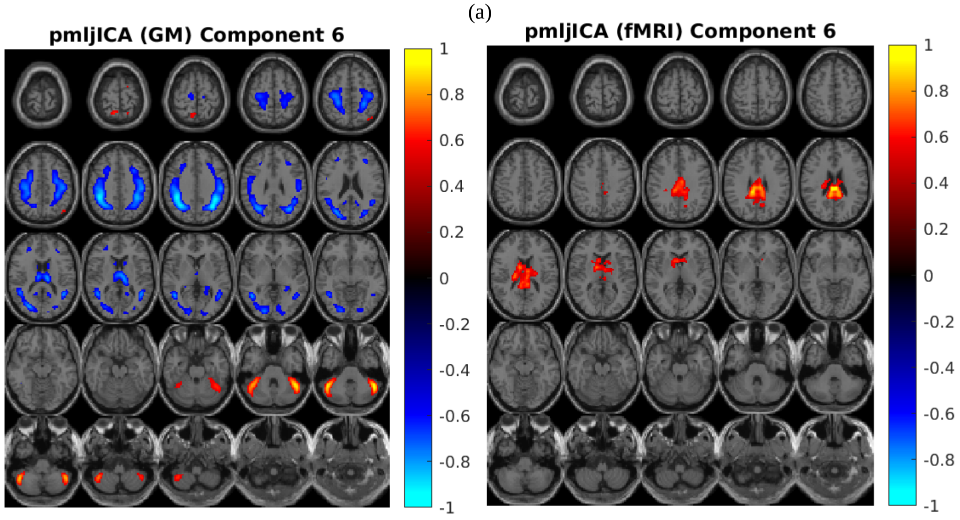


(b)

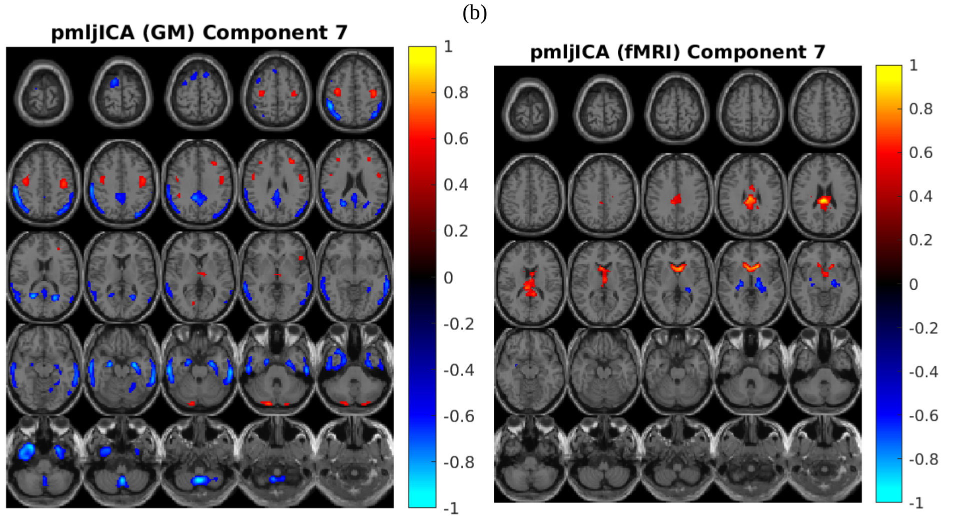


(c)

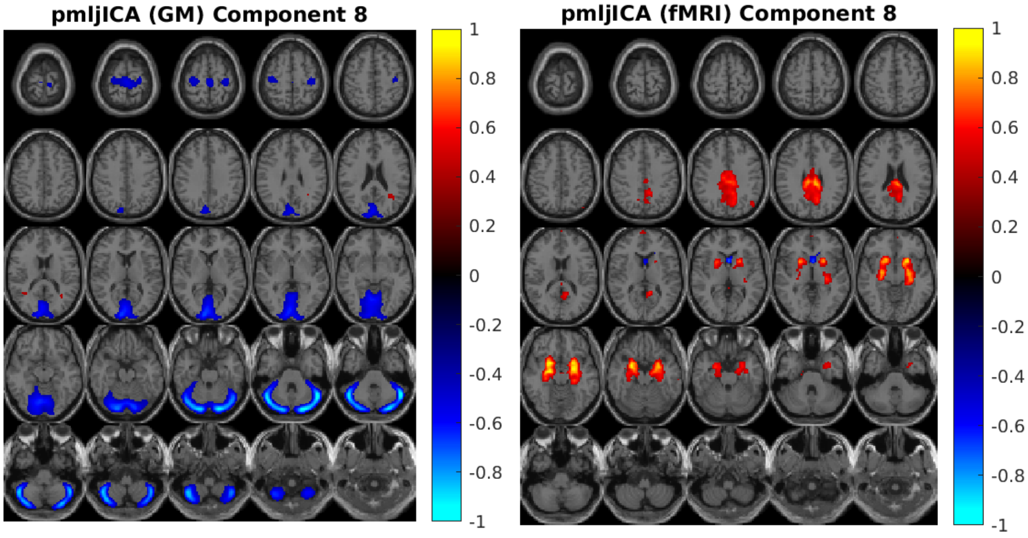
pmljICA (GM) Component 8
pmljICA (fMRI) Component 8
1
0.8
0.6
0.4
0.2
0
-0.2
-0.4
-0.6
-0.8
-1

(d)

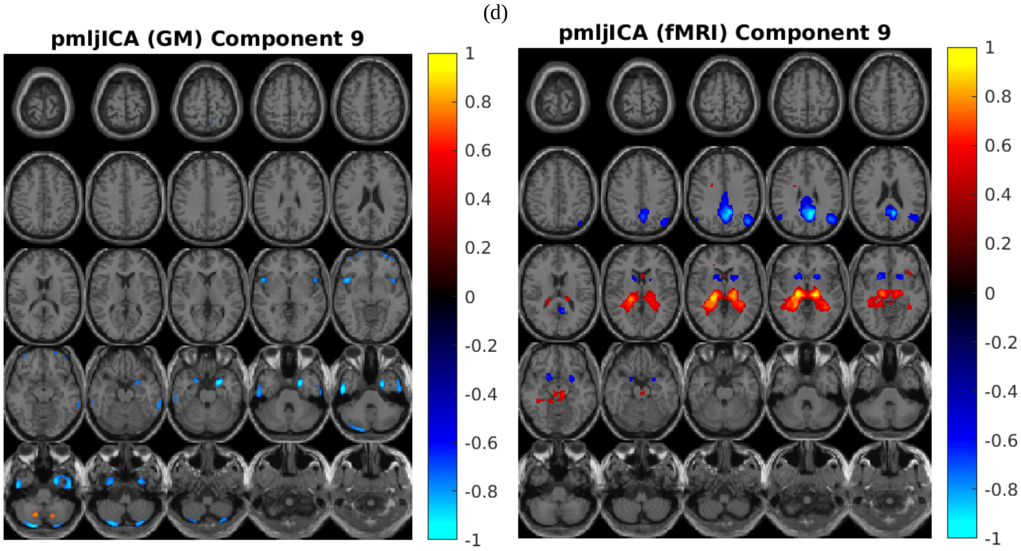
pmljICA (GM) Component 9
pmljICA (fMRI) Component 9
1
0.8
0.6
0.4
0.2
0
-0.2
-0.4
-0.6
-0.8
-1

(e)

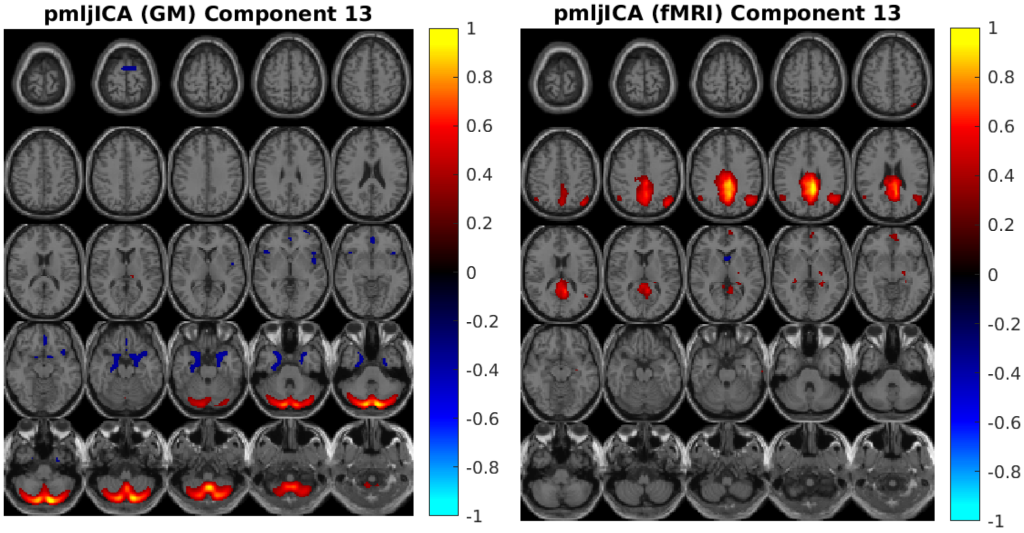
pmljICA (GM) Component 13
pmljICA (fMRI) Component 13
1
0.8
0.6
0.4
0.2
0
-0.2
-0.4
-0.6
-0.8
-1

(f)

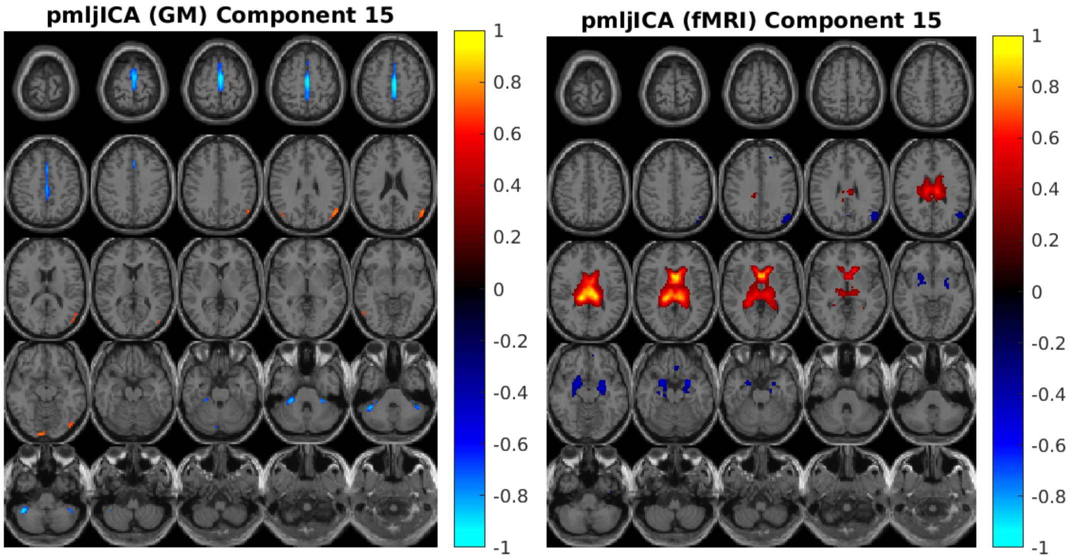


(g)

Figure-1: Joint source map for the Gray matter and rest fMRI. All of activated z score in the source maps are thresholded. The color bar indicates the color mapping of the activated voxels.

The following table summarized brain regions by domain, which are ordered by having volumes greater than 0.4 $cm^3$.

Table-I: Summarized activated brain regions by domain

| **Joint ICs (pmljICA)** | **Brain Regions by Domain** | | |
|---|---|---|---|
| Component 2 | GM | Pos | **AU:** STG(8.3/4.9), TTG(1.2/1.0), MTG(1.1/0.3); **CC:** Insula(7.6/6.1), IFG(3.6/3.6), PHG(2.4/1.6), MeFG(2.2/2.6); **CB:** CL(5.7/6.3), DC(5.4/6.6), CT(2.6/2.8), ISLL(1.5/1.1), Tuber (1.4/1.9), Pyramis (0.9/0.8), Uvula (0.9/0.6); **SC:** Caudate (4.2/4.2), EN(3.5/3.6), TH(2.0/2.0); **DM:** AC(3.5/3.6), Precuneus (2.4/2.6), PC(1.3/2.0); **VI:** FG(3.1/3.8), LG(2.1/4.2); **SM:** PG(1.5/1.6) |
| | | Neg | **AU:** ITG(3.1/3.4), MTG(2.8/3.3), STG(1.0/1.7); **CC:** MeFG(3.6/3.8), SFG(3.0/4.0), MFG (1.7/1.5), Uncus (0.8/2.3); **CB:** ISLL(2.4/4.0), CT(1.3/1.3); **SC:** EN(0.6/1.2), LN(0.5/0.3); **VI:** FG (0.6/1.0) |
| | fMRI | Pos | **CC:** PHG(2.0/1.2); **SC:** EN(1.7/2.2), LN(1.0/1.2); PC(0.1/0.0) |
| | | Neg | **AU:** MTG(0.9/0.0), STG(0.5/0.1); **SC:** Caudate(0.9/0.0), TH(0.1/0.4); **DM:** Precuneus(2.4/0.9), PC(1.2/1.0), AC(0.4/0.0); **VI:** AG(0.6/0.0) |
| Component 6 | GM | Pos | **CB:** CL(4.4/3.1), CT(3.3/3.9), Tuber (2.8/3.4), DC(0.9/0.6); **VI:** FG(0.4/0.1) |
| | | Neg | **AU:** STG(3.3/3.1), MTG(2.6/4.0); **CC:** IPL(4.4/6.7), MFG(3.6/3.3), PHG(0.4/0.0), MeFG(0.3/0.6), IFG(0.2/0.4); **SC:** TH(3.0/2.2), EN(1.9/0.5), Caudate (0.7/0.3); **DM:** Precuneus (3.9/4.9), PC(0.6/0.2); VI: LG(1.7/1.8), AG(0.8/0.7); **SM:** PG(6.5/6.3), PL(0.4/0.3) |
| | fMRI | Pos | **SC:** EN(3.8/6.3), TH(1.5/1.7), Caudate(1.1/2.6), LN(0.1/0.4); **DM:** Precuneus(1.3/0.4), PC(0.9/1.1); |

| | | | |
|---|---|---|---|
| | | Neg | |
| Component 7 | GM | Pos | **CC:** IFG(1.2/0.3), MFG(1.0/0.6); **CB:** Uvula(0.4/0.6); **SC:** TH(0.6/0.2); SM: PG(0.9/0.1) |
| | | Neg | **AU:** MTG(9.1/11.8), ITG(4.5/6.3), STG(1.6/6.6); **CC:** PHG(5.3/1.9), IPL(3.1/8.4), Uncus (2.6/2.8), SFG(1.1/2.6); **CB:** ISLL (1.5/1.2), CL(1.2/0.0), CT(1.0/1.0), DC(0.9/0.0); **SC:** EN (0.6/1.1); **DM:** Precuneus(2.8/1.5), PC(1.2/0.6); **VI:** FG(1.3/1.2), AG(1.3/0.8); **SM:** SPL(1.6/1.7) |
| | fMRI | Pos | **SC:** EN(3.4/3.3), Caudate(1.8/1.3), TH(0.5/0.8); **DM:** PC(0.4/0.4), AC(0.2/0.4) |
| | | Neg | **SC:** EN(2.2/1.4), TH(1.5/0.1), LN(0.3/0.4) |
| Component 8 | GM | Pos | |
| | | Neg | **CC:** MeFG(1.1/1.4); **CB:** Tuber(7.1/6.7), CL(6.4/7.5), CT(5.4/5.8), ISLL(3.7/4.7), Pyramis (2.0/2.0), DC(10.9/11.0), Cuneus (10.2/15.4), Uvula (1.2/1.0); **DM:** PC(0.4/1.0), Precuneus(0.3/2.6); **VI:** LG(7.5/13.4), FG(2.2/4.2); **SM:** PG(0.6/0.3); |
| | fMRI | Pos | **CC:** PHG(7.5/5.4), Uncus (1.8/1.4), IFG(0.4/0.8); **SC:** EN(7.3/4.6), LN(5.5/4.7); **DM:** Precuneus(4.9/2.0), PC(3.1/1.0); **VI:** AG(0.5/0.0); |
| | | Neg | **EN:** EN(0.4/0.2) |
| Component 9 | GM | Pos | |
| | | Neg | **AU:** ITG(1.2/1.3), MTG(1.2/1.3), STG(0.6/1.2); **CC:** Uncus(2.6/0.6), IFG(0.4/0.6), MFG(0.4/0.3), SFG(0.4/0.3); **CB:** ISLL(0.8/1.0), Pyramis (0.4/0.6), Tuber (0.2/0.7); **VI:** FG(0.2/0.4) |
| | fMRI | Pos | **SC:** EN(5.6/6.6), TH(5.4/5.5), LN(0.4/0.4) |
| | | Neg | **AU:** MTG(1.2/0.1), STG(0.8/0.2); **CC:** IPL(1.3/0.0), PHG(0.4/0.4); **SC:** LN(1.9/1.7), EN(1.1/1.7), Caudate (0.3/0.7); **DM:** Precuneus (6.8/2.8), PC(1.9/0.8); **VI:** AG(2.6/0.3) |
| Component 13 | GM | Pos | **CB:** ISLL(6.6/6.8), DC(3.9/5.4), Pyramis (3.2/3.1), Tuber (2.9/3.5), Uvula (2.4/2.3), CT(1.7/1.5), Pyramis of Vermis (0.4/0.2); **VI:** FG(0.1/0.4) |
| | | Neg | **AU:** STG(1.8/1.1); **CC:** Uncus (2.2/3.3), IFG(1.9/1.2), PHG(1.7/4.4), SFG(0.7/0.1), MeFG(0.5/0.3), Insula (0.5/0.0); |
| | fMRI | Pos | **AU:** STG(1.7/0.3), MTG(1.3/0.1); **CC:** IPL(2.4/0.7), MeFG(1.3/0.2); **SC:** EN(4.0/3.0), TH(0.2/0.0); **DM:** PC(4.6/3.5), Precuneus (10.4/6.7); **VI:** AG(3.0/0.6) |
| | | Neg | |
| Component 15 | GM | Pos | **AU:** MTG(1.0/0.0); **VI:** AG(0.7/0.3) |
| | | Neg | **CC:** MeFG(2.4/1.9), SFG (0.6/0.5); **CB:** CL(0.6/2.0), CT(0.3/1.2); **SM:** PL(2.5/1.2) |
| | fMRI | Pos | **CC:** PHG(0.5/0.3); **SC:** TH(8.3/8.8), Caudate (5.4/4.2), EN(14.8/16.1); |
| | | Neg | **AU:** MTG(1.1/0.0); **CC:** PHG(2.6/2.5), Uncus (0.3/0.6); **SC:** EN(1.1/0.9), LN(0.4/0.7); **VI:** AG(1.6/0.0) |

Brain Domains

**AU:** Auditory Domain
**SC:** Sub-cortical Domain
**SM:** Sensorimotor Domain
**VI:** Visual Domain
**CC:** Cognitive-control Domain
**DM:** Default-mode Domain
**CB:** Cerebellar Domain

Brain Regions

**STG:** Superior Temporal Gyrus
**TTG:** Transverse Temporal Gyrus
**MTG:** Middle Temporal Gyrus
**ITG:** Inferior Temporal Gyrus
**IFG:** Inferior Frontal Gyrus
**PHG:** Parahippocampal Gyrus
**MeFG:** Medial Frontal Gyrus
**SFG:** Superior Frontal Gyrus
**MFG:** Middle Frontal Gyrus
**ISLL:** Inferior Semi-Lunar Lobule
**SPL:** Superior Parietal Lobule
**IPL:** Inferior Parietal Lobule
**CT:** Cerebellar Tonsil

**AG:** Angular Gyrus
**LG:** Lingual Gyrus
**PG:** Postcentral Gyrus
**PL:** Paracentral Lobule
**EN:** Extra-Nuclear
**LN:** Lentiform Nucleus
**FG:** Fusiform Gyrus
**PC:** Posterior Cingulate
**CL:** Culmen
**DC:** Declive

Pos – Positive Activity
Neg – Negative Activity
